# Learning of Spatial Properties of a Large-scale Virtual City with an Interactive Map

**DOI:** 10.1101/539080

**Authors:** Sabine U. König, Viviane Kakerbeck, Debora Nolte, Laura Duesberg, Nicolas Kuske, Peter König

## Abstract

On the basis of embodied/-enacted theories of the mind, investigations of spatial cognition related to real world environments have become current research interests. How this perspective relates to acquiring spatial knowledge not by active exploration, but through map learning, however, remains unresolved. Therefore, we designed a large virtual city comprised of over 200 houses, suitable for active exploration or spatial learning based on a map. Here, we report our results after single and repeated 30-minute training sessions using an interactive city map. We tested subjects’ knowledge of the orientation of houses towards cardinal north and of two houses relative to each other and the locations of two houses towards each other in a pointing task. Our results revealed that a single training session was sufficient to repeatedly view the majority of houses covering a large area of the city. However, repeated training sessions were necessary to improve the performance level and reveal significant differences between tasks. In contrast to a previous study in a real world city, performance in the two orientation tasks was better than in the pointing task. The lack of distance and the lack of angular difference effects onto task performance suggest the use of a global reference frame. Performance was positively correlated with a self-report on spatial abilities (FRS) in the absolute orientation and pointing task but not the relative orientation task. Overall, our results suggest that training with an interactive city map enhances abstract knowledge, not directly available from an embodied perspective.

## Introduction

According to theories of embodied/enacted cognition spatial navigation unfolds in the interaction of the navigator with his real world surroundings (Engel, Maye, Kurthen, & König, 2013; O’Regan & Noe, 2001). In known environments, spatial navigation, e.g. to get to the nearest bakery or go home after work, is based on memorized spatial knowledge and bodily movement in the environment. Bodily movement provides the information about multimodal motor, proprioceptive and vestibular changes, which are important for spatial learning (Grant & Magee, 1998; Riecke et al., 2010; Ruddle, Volkova, & Bülthoff, 2011; Ruddle, Volkova, Mohler, & Bülthoff, 2011; Waller, Loomis, & Haun, 2004). Interaction with the environment is a crucial part of learning in spatial navigation, thus becoming more and more important for spatial navigation research (Gramann, 2013).

While navigating in an unknown surrounding, people acquire spatial knowledge through direct experience. They develop knowledge of landmarks, i.e. salient objects in the environment, route knowledge, i.e. connecting routes between landmarks, and through the integration of acquired route knowledge so called survey knowledge, a map like representation of metric spatial relationships (Siegel & White, 1975). These three types of spatial knowledge can be learned after only one exposure to the environment, but there are large individual differences in spatial knowledge acquisition (Ishikawa & Montello, 2006; Montello, 1998; Wiener, Büchner, & Hölscher, 2009). Empirical studies showed that early spatial learning directly includes knowledge about metric properties of the environment like distances and angles between locations (Klatzky et al., 1990; Montello & Pick, 1993). With increasing familiarity of a location, spatial knowledge becomes more precise and certain (Ishikawa & Montello, 2006; Montello, 1998). Spatial learning through active navigation is supposed to especially improve route knowledge (Meilinger, Frankenstein, & Bülthoff, 2013; Shelton & McNamara, 2004; Taylor, Naylor, & Chechile, 1999; Thorndyke & Hayes-Roth, 1982). However, the kind of spatial knowledge that is used depends on the respective task and likely combines multiple types of knowledge (Meilinger et al., 2013).

Beside embodied experience, maps are also important for spatial learning (Meilinger et al., 2013; Shelton & McNamara, 2004). In western countries cartographical 2-dimensional (2D), north-up maps from a bird’s eye view are most often used. Those maps depict the real world in downscale while keeping the proportions intact. Thus, distances and locations as well as spatial orientation resemble the real world situation. Therefore, cartographical maps are supposed to support acquisition of survey knowledge (Meilinger et al., 2013; Taylor et al., 1999; Thorndyke & Hayes-Roth, 1982). This is also supported by previous research that reported better spatial orientation in a pointing task when participants faced north in a familiar environment (Frankenstein, Mohler, Bülthoff, & Meilinger, 2012). They argued that the improved performance with the northern orientation makes it likely that participants acquired their knowledge with a north-up map as research found that pointing accuracy improved when tested and learned viewpoints were aligned (Shelton & McNamara, 1997). In modern times, technology allows to switch from cartographical 2D maps to 3-dimensional (3D) interactive maps like Google Street View on digital devices. In interactive city maps, observers change their perspective from a bird’s eye view to a pedestrian view, mimicking an embodied perspective, while navigating. A recent report compared the performance in orientation tasks after learning with 2D and 3D city maps (Oulasvirta, Estlander, & Nurminen, 2009). They found that learning from 2D maps yielded better performance than from 3D maps, even in participants who were used to 3D navigation through gaming experience. Overall, 3D maps supply a pedestrian perspective of the environment, whereas 2D maps provide the observer with the information of metric properties of large-scale environments thus supporting different aspects of spatial knowledge acquisition.

The source of spatial learning also influences the kind of reference frame, an important aspect in spatial cognition, in which spatial features are learned and memorized. The most important distinction of spatial reference frames is drawn between egocentric and allocentric reference frames (Burgess, 2006; Klatzky, 1998; Mou, McNamara, Valiquette, & Rump, 2004). An egocentric reference frame relates the environment to the physical body of the navigator (Klatzky, 1998). Meilinger et al. (Meilinger et al., 2013) found that route knowledge improved when participants responded in a route recall task from a pedestrian perspective. This suggests that spatial knowledge learned by active navigation is coded in an egocentric reference frame (Shelton & McNamara, 2001). Also spatial updating, which is caused by changes in somatosensory information, is coded in an egocentric reference frame (Riecke, Cunningham, & Bülthoff, 2007; Wang & Brockmole, 2003). Instead, spatial knowledge coded in an allocentric reference frame is based on allocentric bearing and distance independent of the physical body of the observer (Klatzky, 1998). Thus, getting acquainted with an environment by a map, which provides information of cardinal directions and spatial orientation presumably supports spatial knowledge coded in an allocentric reference frame. Even though spatial knowledge is stored in distinct reference frames, they are supposed to develop together (Nardini, Burgess, Breckenridge, & Atkinson, 2006) and to be combined for active navigation (Burgess, 2006; Gramann, 2013; Ishikawa & Montello, 2006; Meilinger et al., 2013).

Spatial knowledge that is coded in an egocentric reference frame can easily be understood within an action-oriented approach. But can spatial properties coded in an allocentric reference frame be learned through active navigation as well? Testing pointing accuracy in a large-scale environment that was learned by active navigation, McNamara, Rump, and Werner (2003) found an improved pointing accuracy when tested objects were aligned with salient aspects of the environment, which serve as the reference in an allocentric frame. Also Brunyé et al. (2015) found that cognitive maps can preferably be oriented towards salient environmental features, thus in allocentric terms, when the environment was learned by active navigation. In cases where the salient environmental features were aligned with north, pointing tasks revealed the best performance when these features were aligned to the north direction (Brunyé et al., 2015; Marchette, Yerramsetti, Burns, & Shelton, 2011). A previous study (König, Goeke, Meilinger, & König, 2017) investigated allocentric spatial knowledge retrieval after everyday navigation in the home-town of Osnabrück, Germany. They used photographs of houses and streets as stimuli and tested spatial knowledge of orientation of houses and streets towards cardinal north, relative orientation of two houses and of two streets, respectively, and relative location of two houses in a pointing task. Investigating spontaneous knowledge retrieval, houses were best remembered in house to house relations, whereas streets were preferentially coded in relation to cardinal directions. Time for cognitive reasoning also improved the knowledge of cardinal orientation of houses. This type of spatial knowledge obtained by everyday navigation in one’s hometown is achieved by a combination of active navigation and the use of maps. In these circumstances, it is difficult to resolve which spatial properties are learned from one or the other source.

In the presented study, we therefore explored knowledge acquisition of spatial properties in a large virtual city with controlled exploration, reporting here the learning with an interactive city map. Progress of technology in virtual reality (VR) renders it possible to design environments in VR that support the investigation of spatial navigation and spatial learning with respect to embodied navigation and the possibility to investigate spatial learning under controlled conditions. For this reason, we built a large virtual city named Seahaven, which would cover 500m x 431m in real world measurements and contains 214 houses. In a separate study, we report the results after training in VR. In the present study, all participants explored the virtual city by means of an interactive city map. This map displayed the front-on screenshots of houses when the participant clicked on the respective house in the top-view north-up map. Additionally, we introduced a pre-training task consisting of one example trial for each of the task conditions with house stimuli of a real-world city to familiarize participants with the tested spatial properties and allow that they were aware of the nature of later tests. To test the acquired spatial knowledge, we adjusted and performed appropriately adjusted versions of the three navigation tasks, which were used in the previous study (König et al., 2017). The absolute orientation task investigates the knowledge of orientation of a single house towards north. The relative orientation task investigates the knowledge of orientation of two houses towards each other. The pointing task investigates the knowledge of relative location of two houses to each other. We used front-on screenshots of houses of the virtual city as task stimuli and tested all tasks with spontaneous response (within 3 seconds) and time for cognitive reasoning (infinite time to respond). Taken together, we expected that with infinite response time for cognitive reasoning, participants would perform better than when spontaneous decisions in only 3 seconds were required. As participants got to know the virtual city with an interactive city map that provides metric spatial properties, e.g. of cardinal directions, orientation and distance, we expected that participants would perform better at determining the absolute orientation of a single house towards north than the relative orientation of two houses or the relative location of two houses to each other. Further, we investigated familiarity of tested houses, distance and angular difference between stimuli, subjectively rated spatial ability (FRS) and increased training time as possible factors that might have an influence on the task performance. Overall, we investigated controlled spatial learning with an interactive city map of a virtual city to get more insight into learning of spatial properties and examine whether these are comparable to knowledge of spatial properties learned in a real city (König et al., 2017).

## Methods

### Objective and procedure

In this study, we aimed at investigating whether free spatial exploration of an unknown virtual city with an interactive map would yield comparable results in spatial knowledge to that acquired in a real world city by everyday navigation (König et al., 2017). The sequence of methods is in close analogy to that previous study (see below). All participants started with a response training to familiarize them with the choice of the directional arrows on the stimuli and the 3 seconds response mode (König et al., 2017). Further, they underwent additional task instructions and performed a pre-task training to sensitize them to pay attention to important spatial information during their training to solve the post-training tasks. We investigated acquisition of spatial knowledge of the virtual city Seahaven by three different means. We report here the results of participants, who were instructed to use an interactive city map of Seahaven. Other groups of participants explored the city in VR with or without the feelSpace belt, a device supplying information about magnetic north (Kaspar et al., 2014; König et al., 2016). These data are, however, not covered in the present report. After the exploration, all participants were tested on three different tasks. These spatial tasks were designed as two-alternative forced-choice tasks (König et al., 2017) and measured the knowledge of the absolute orientation of a single house towards the north cardinal direction, the relative orientation between two houses, and pointing from the location of one house to the other (see below). Each task type was performed in two time conditions: with response time restricted to 3 seconds for spontaneous decisions and infinite response time for cognitive reasoning. At the end of the experiment, all participants filled in the FRS questionnaire, a self-report on spatial abilities (Münzer & Hölscher, 2011).

### Participants

77 young healthy adults (40 females, mean age of 24,0 years, SD = 3,9) took part in our study using an interactive map for their training. Seven participants had to be excluded because of technical problems during the first measurement. Out of the remaining 70 participants 22 trained and performed all tests repeatedly on three separate days (11 females, mean age of 23,8 years, SD = 3,1). Before the start of the experiment, participants were informed about the purpose and the procedures of the experiment and gave written informed consent. Each participant was either reimbursed with nine Euros per hour or earned an equivalent amount of ‘‘participant hours,’’ which are a requirement in most students’ study programs. Overall, the experiment took about two hours each session. The study was approved by the ethics committee of the Osnabrück University in accordance with the ethical standards of the Institutional and National Research Committees.

### Pre-task information and - training

Each participant underwent task instructions and task training before the start of the free exploration time of Seahaven. We included this pre-task information, because a pilot study revealed that during their free exploration of the city participants sometimes focused more on aspects, e.g. house design, not relevant for the spatial tasks. The instructions were given by written and verbal explanations using photographs of houses in the city of Osnabrück that were used as stimuli in a previous study (König et al., 2017). Participants then performed a pre-task training with one example of all post-training tasks each in both time conditions to gain a better insight into the actual task requirements. The stimuli in the pre-task training were again photographs of houses of the city Osnabrück to avoid possible transfer of training effects onto the stimulus set of Seahaven. Except for the stimuli, the pre-tasks resembled exactly the post-training tasks design (see in experimental design below). We performed the pre-task training to familiarize participants with the spatial knowledge that was tested in the post-training tasks.

### The Virtual City „Seahaven“

To be able to investigate the spatial exploration and acquisition of spatial knowledge of an unknown environment, we designed a virtual city, called Seahaven. The name Seahaven is derived from the name of the town in Peter Weir’s 1998 film ‘The Truman Show’, in which the main character Truman Burbank is also living in a small town with no possibility to leave while every moment is watched and analyzed by many people, albeit in the form of a reality TV show. Our virtual city Seahaven was built in the Unity® game engine. Seahaven covers 500×431 unity units. 1 unity unit was designed to resemble 1 m in real world measure. Thus, Seahaven covers 0,216 km^2^ in real world measures (Fig. 1). Seahaven contains 214 houses in diverse styles. The number of houses sharing a certain orientation towards North is approximately equally distributed in steps of 30° (0° to 330°). Compared to many virtual environments used for spatial navigation research, Seahaven is large and complex. Compared to the real city of Osnabrück, the size of Seahaven is approximately one fifth of the inner city surrounded by the “Wall” (Fig. 1 B). This is a size that can be well explored within half an hour. For the purpose of the following spatial navigation tasks, Seahaven does not contain tall landmarks or specific city districts and the street system does not follow an ordered grid (i.e. “Manhattan style”). All files required for the conduction of experiments are available on the GitHub account of one of the authors (V. Kakerbeck).

**Fig. 1:**
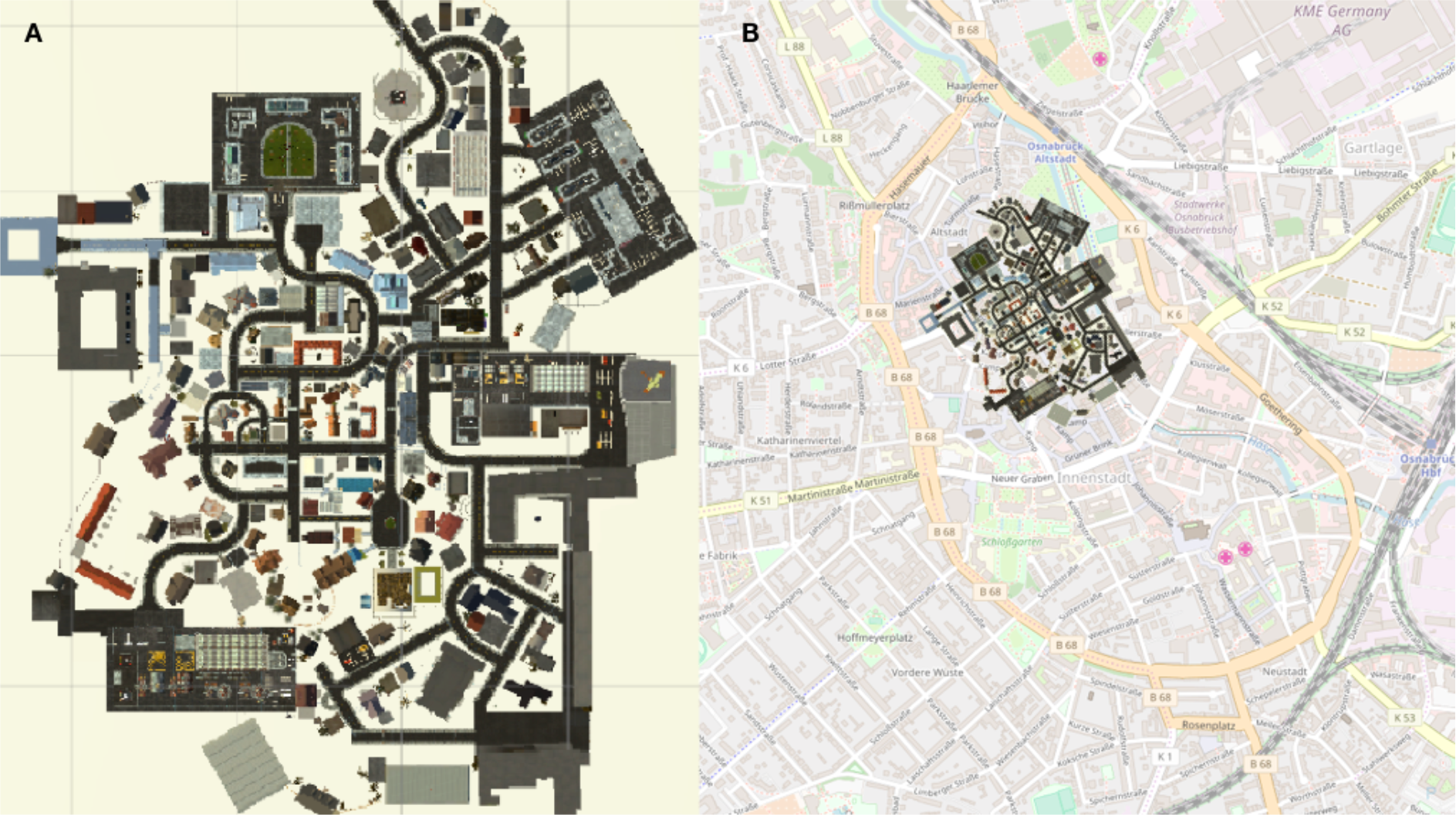
(A) City map of Seahaven depicting all houses that can be clicked on, to display a screenshot of the front-on view of the respective house (B) City map of the virtual city Seahaven overlaid onto the city map of central Osnabrück.

### Stimuli

Our stimuli were front-on screenshots of 193 houses of the overall 214 houses in Seahaven (examples Fig. 2 to 5). We could not use all houses because the design of the city did not allow every house to be viewed from an angle that could be used to take the screenshot. Furthermore, a few houses looked too similar to each other and therefore had to be excluded as well. We took the screenshots in the virtual environment from a viewpoint that would resemble approximately 5-meter distance to the corresponding house. Specifically, the screenshots were taken from a position on a street that could be walked past by the participants during the VR city exploration. The screenshots were scaled to 2160 × 1920 pixels, so that each screenshot was presented in full screen on one monitor. For the prime stimuli in the relative orientation and pointing tasks we used the screenshots of 18 houses that were most often viewed in a VR pilot study (although the present subjects experienced the city exclusively by the interactive map). Thus, we aimed to ensure that the prime stimuli were well known to most, if not all participants. These prime houses were distributed equally over the possible angles differing from VR cardinal north in steps of 30° and were distributed well across the city. In the post-training tasks we used a subset of all screenshots that were used in the interactive city map training.

**Fig. 2:**
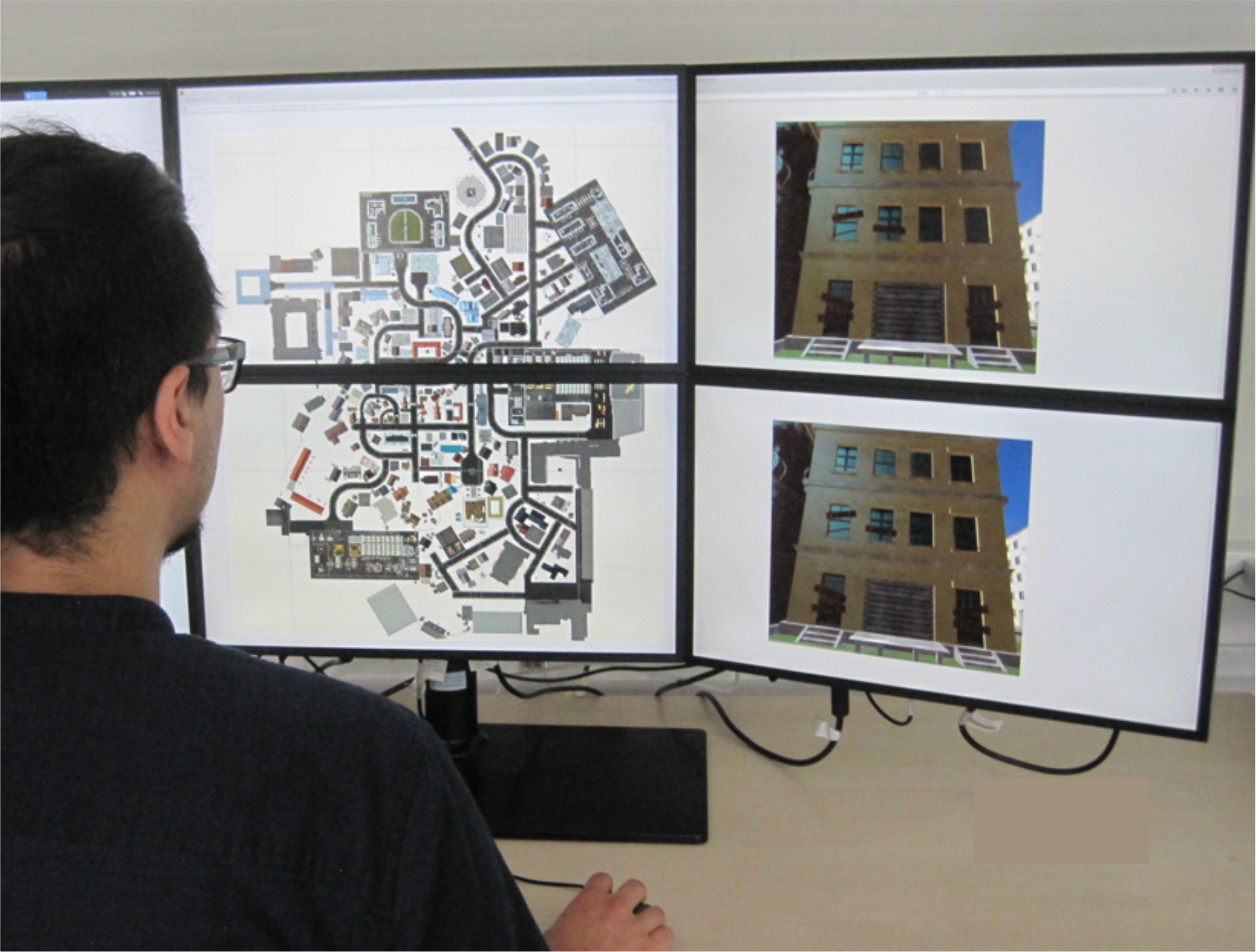
Experimental Set-up of the interactive city map of Seahaven. Left: Birds-eye view, north up city map. Right: The Screenshot of the clicked house appears on the two-stacked screens on the right.

### Training in Seahaven with an Interactive Map

To get to know the city of Seahaven, participants trained in single and repeated sessions of 30 minutes with a two-dimensional interactive city map. Participants sat approximately 60 cm from a six-screen monitor setup in a 2 × 3 screen arrangement. During the training, the map was presented on the two central screens, one above the other (Fig. 2, left). This map resembled a traditional city map with a north-up orientation and a bird’s-eye view. It was implemented using html, jQuery and CSS. The map provided the participants with task relevant spatial information about the location, orientation and relation of houses. By adding an interactive component, participants were also provided with the screenshots of front-on views of 193 houses in Seahaven that were also used as post-training task stimuli. In order to view the houses’ screenshots, participants moved with a mouse over the map. When hovering over a house, a red dot appeared on one side of the respective house. The red dot indicated the side of the house that was displayed in the screenshot. By clicking on this house the screenshot was displayed twice on the two right screens of the monitor (the same image above each other) (Fig. 2, right). How often houses were clicked on was recorded and later used to determine participants’ familiarity with the stimuli and which houses were looked at to investigate the part of Seahaven that participants visited. The two screens on the left side were not used in training or testing. During the training participants got feedback on how much time had passed after 15 and 25 minutes. They had a short break of 2 to 5 minutes between map training and testing of post-training tasks. In summary, the interactive map provided participants with a city map of Seahaven and the front on views of the majority of houses of Seahaven.

### Post-training Task Design

The post-training tasks have been described in a previous study by König et al. (König et al., 2017) in detail. Briefly here: During the post-training tasks, the images were presented on the two central screens, while the other four monitors to the sides were switched off. Written instructions were given on the screen. Participants responded by pressing a button on a control box.

#### Absolute orientation task

In the absolute orientation task, we measured participants’ knowledge about the orientation of single houses in relation to the cardinal direction north. Therefore, we designed a two-alternative forced-choice (2AFC) task with front-on views of houses of the virtual city Seahaven as stimuli (Fig. 3). We presented the same house stimulus on the stacked middle screens of our 6-screen monitor (Fig. 3C). An arrow within an ellipsoid depicting a compass was overlaid on each of the two stimuli. One of the arrows pointed correctly to the northern direction, whereas the other arrow pointed randomly in a direction that diverged from north by some amount between 0° and 330° in steps of 30°. Participants had to choose, which arrow pointed correctly towards north by pressing either the “up” (upper screen) or “down” (lower screen) button on the response box. We used two different time conditions: 3 seconds and infinite time to respond to investigate the influence of spontaneous decisions compared to decisions with time for cognitive reasoning, respectively. In case the subject did not respond within the time limit, the trial was counted as incorrect. This was done as well in the other tasks. Both time conditions contained 36 trials. One trial consisted of a grey screen shown for 5 seconds on both middle screens, which was followed by the stimulus set. After the press of a response button, either within 3 seconds or with infinite time for the decision, again the two central grey screens appeared followed by another stimulus set. The gray screens were interposed between the experimental trials to have the same time course as in the two other tasks (see absolute orientation and pointing task). The two time conditions were blocked and introduced by written instructions on the screens.

**Fig. 3:**
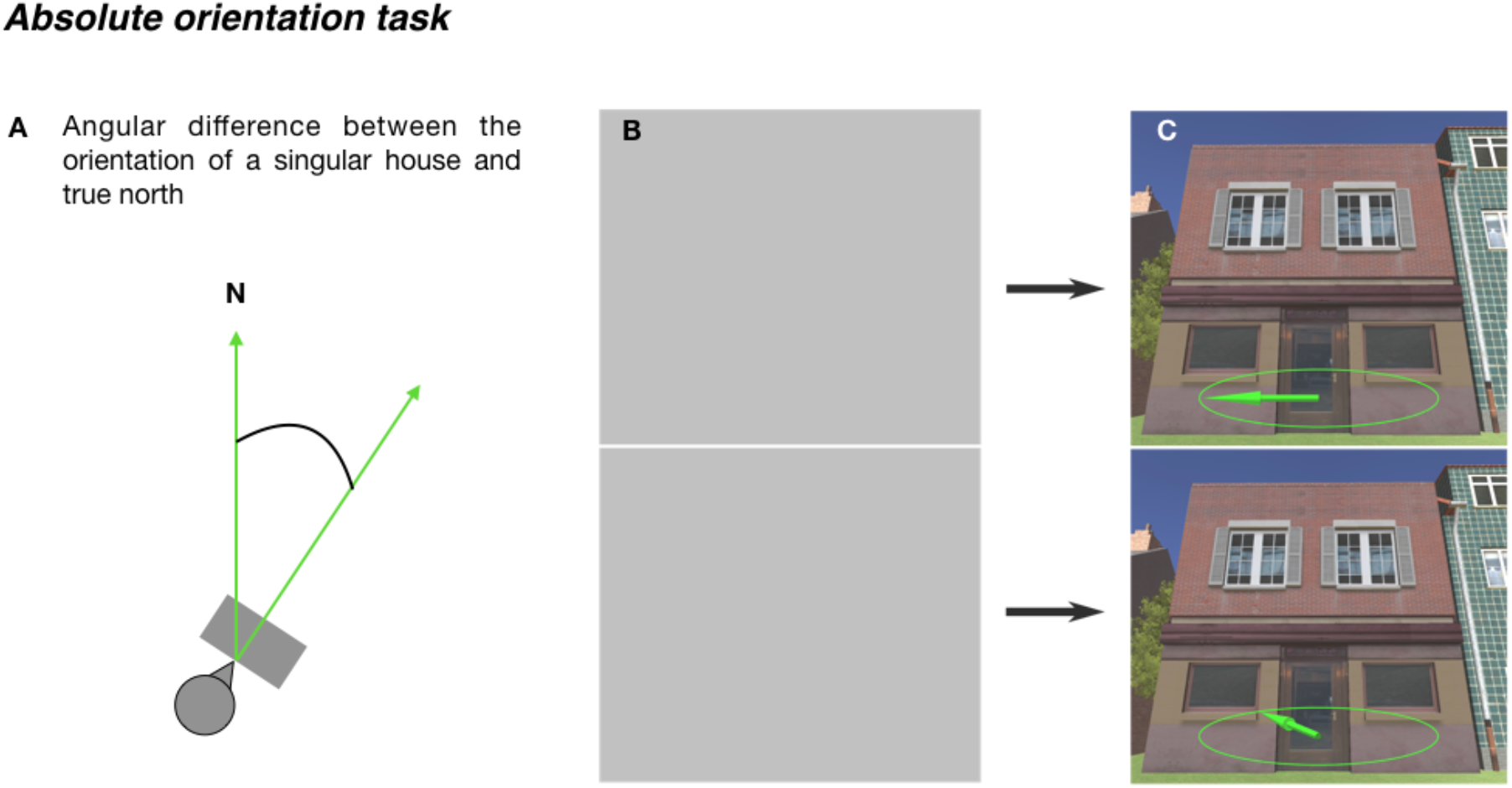
Absolute orientation task. (A) Schema of the absolute orientation task. (B and (C) The illustration shows one sample trial of the absolute orientation task. (B) Depicts a grey screen shown for 5 seconds to fit the experimental condition in the other tasks. It is followed by a stimulus (C) using a screenshot of a house in the virtual city shown until button press max for 3 seconds or for infinite time until button press on two monitors, one above the other. The images were overlain with arrows within an ellipsoid, depicting a compass. In each example, one arrow points correctly to north and the other randomly to a different direction. Participants had to choose the arrow that correctly pointed towards north.

#### Relative orientation task

In the relative orientation task, we measured the knowledge to estimate the orientation of houses relative to each other (Fig. 4). We then designed a 2AFC task with front-on views of houses of the virtual city Seahaven as stimuli again with either 3 seconds or infinite time to respond. The general task design resembled the one of the absolute orientation task, but here each trial depicted a stimulus set consisting of a fixed triplet of images: one priming image and two target images (Fig. 4 B and C). In each trial, the priming image was first shown on both middle screens for 5 seconds. After the priming stimulus was turned off, the two target stimuli appeared, one on the upper screen and the other on the lower screen. For a few houses it was not possible to perfectly align prime and target stimuli, the task of the participants was to select the target image that was more closely aligned (+/− 10°) with the orientation of the priming image by pressing either the “up” or “down” button on a response box, respectively. Again, this sequence was repeated with 36 stimulus sets in each time condition and the time conditions were blocked and introduced by written instructions on the screens.

**Fig. 4:**
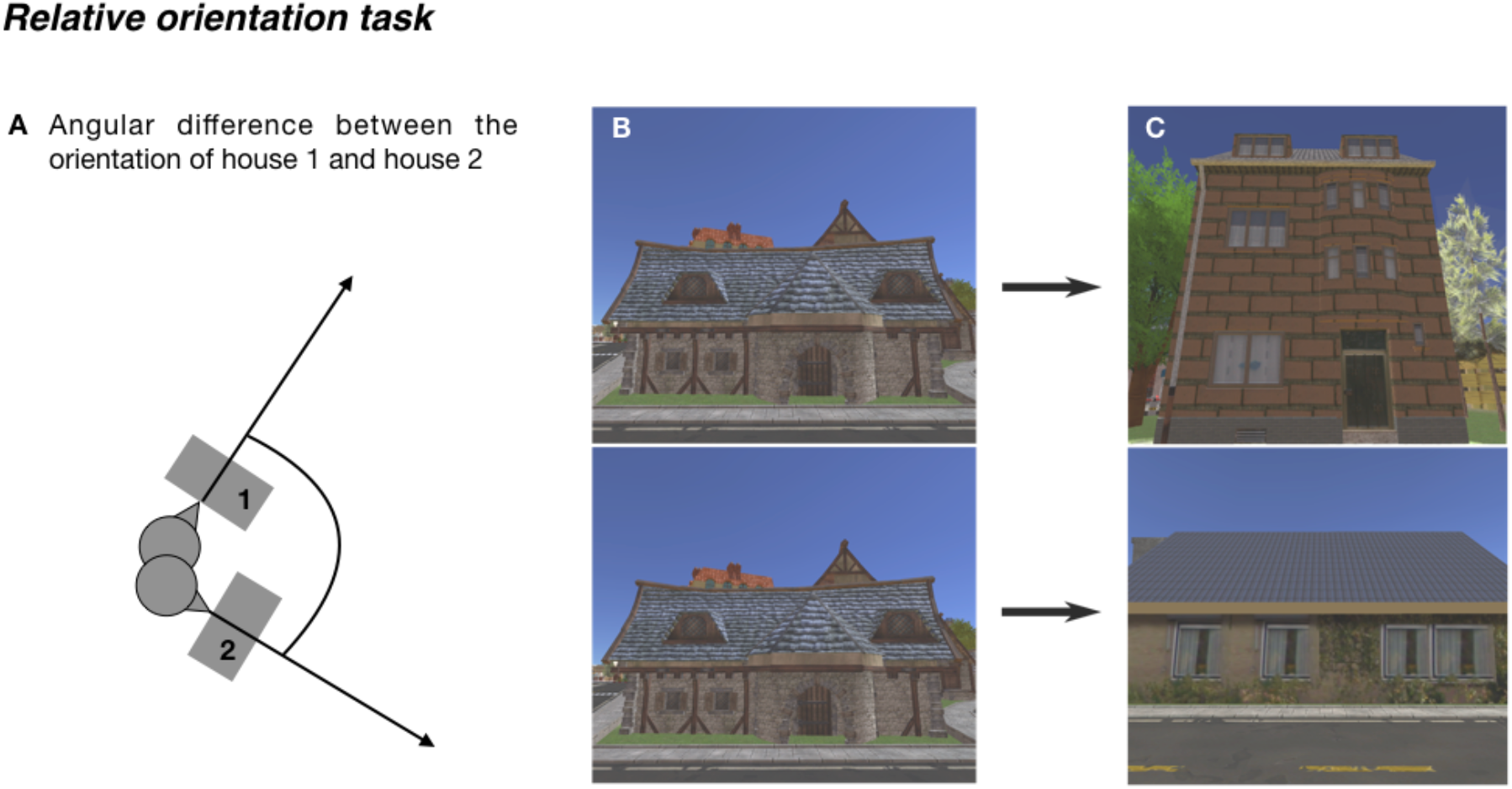
Relative orientation task. (A) Schema of the relative orientation task. (B and C) The illustration shows one sample trial of the relative orientation task. (B) First, a priming image (screenshot from a house in Seahaven) was shown for 5 seconds. (C) Then, two different target images (screenshots from houses in Seahaven) appeared on the two screens, one above the other. One of the target images had the same orientation as the priming image (within a range of +/− 10° compared to the prime image), while the other differs by varying degrees. The task of the participants was to select the image that was oriented in the same way as the priming image.

#### Pointing task

In the pointing task, an established paradigm in spatial navigation studies, we investigated the knowledge of the spatial relation between the locations of two houses in Seahaven (Fig. 5). We used the same stimulus material of houses as before. Here, we defined pairs of houses with a priming stimulus and a target stimulus. First, a priming image appeared on two stacked screens for 5 seconds (Fig. 5B), which was followed by a target stimulus, the same on both screens, either for 3 seconds or until a response. Similar to the absolute orientation task, we overlaid the target image with an arrow within an ellipsoid, this time depicting the pointing direction. On one screen, the arrow was correctly pointing into the direction from the target to the priming image location. On the other screen, the arrow was pointing in a random direction differing with varying degrees (30° to 330° in 30° steps) from the correct direction (Fig. 5C). In the 2AFC task, participants had to choose the target stimulus on which the arrow pointed correctly from the target stimulus towards the priming stimulus by pressing either the “up” or “down” key. The pointing task consisted again of 36 trials in each time condition, which were blocked and introduced by written instructions on the screen.

**Fig. 5:**
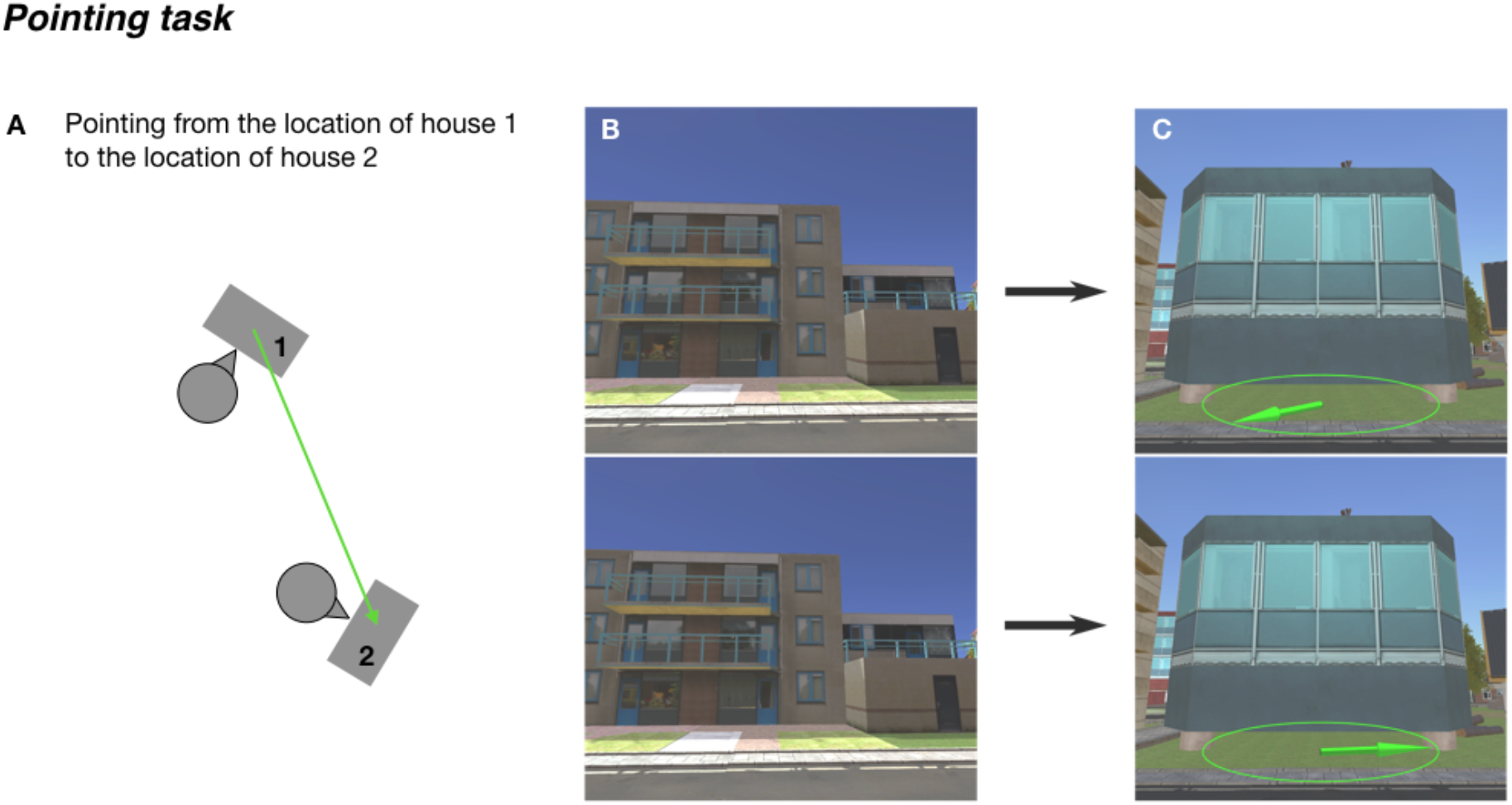
Pointing task. (A) Schema of the pointing task. (B and C) The illustration shows one sample trial of the pointing task. (B) First, a priming image (screenshot from a house in Seahaven) was shown for 5 seconds on both monitors. (C) Then, a target image (screenshot from a house in Seahaven) appeared on the two screens. One of the target images was overlaid with an arrow pointing correctly towards the priming image, whereas the other was overlain with an arrow pointing into a direction that differed by varying degrees. The participant had to choose the arrow that pointed correctly towards the priming image.

### Randomization

As we expected larger angular differences to be easier to learn, we used houses of Seahaven from 0° to north in steps of 30° up to 330° in equal proportions. In the post-training tasks, we randomized the angular difference in the absolute orientation task between the correct arrow and the wrong arrow (Fig. 3c), in the relative orientation task between the two target houses (Fig. 4c), and in the pointing task between the correct and wrong arrow of the target house pointing to the prime house (Fig 5c). Also, the differences of target and distractor angles were equally distributed. We randomized the blocks of task and time condition to prevent a systematic bias of learning effects over time. In summary, to prevent bias effects, we randomized our design in all relevant aspects.

### FRS Questionnaire

At the end of the measurements, participants filled in the FRS questionnaire (“Fragebogen Räumlicher Strategien”, translated “Questionnaire of Spatial Strategies (Münzer & Hölscher, 2011)) to get insight into preferred spatial strategies and abilities. The FRS evaluates three different scales: the global-egocentric -, survey - and cardinal directions scale.

## Results

We first present results after a single training session, in case of repeated training only the first session. An analysis of the repeated sessions is described further below.

### Training results

We determined how many houses the participants looked at and how often these houses were clicked on. The number of houses that were looked at by each participant ranged from 102 (52,85%) to 188 houses (97,41%) out of 193 possible houses, resulting in a grand average of 161 houses (83,42%) (Fig. 6A). The mean number of clicks on a house made by each participant ranged from 0,67 to 8,14 clicks, resulting in a grand average of 3,95 clicks (Fig. 6B). Thus, the half hour training time was sufficient to consistently explore a larger fraction of all houses.

**Fig. 6:**
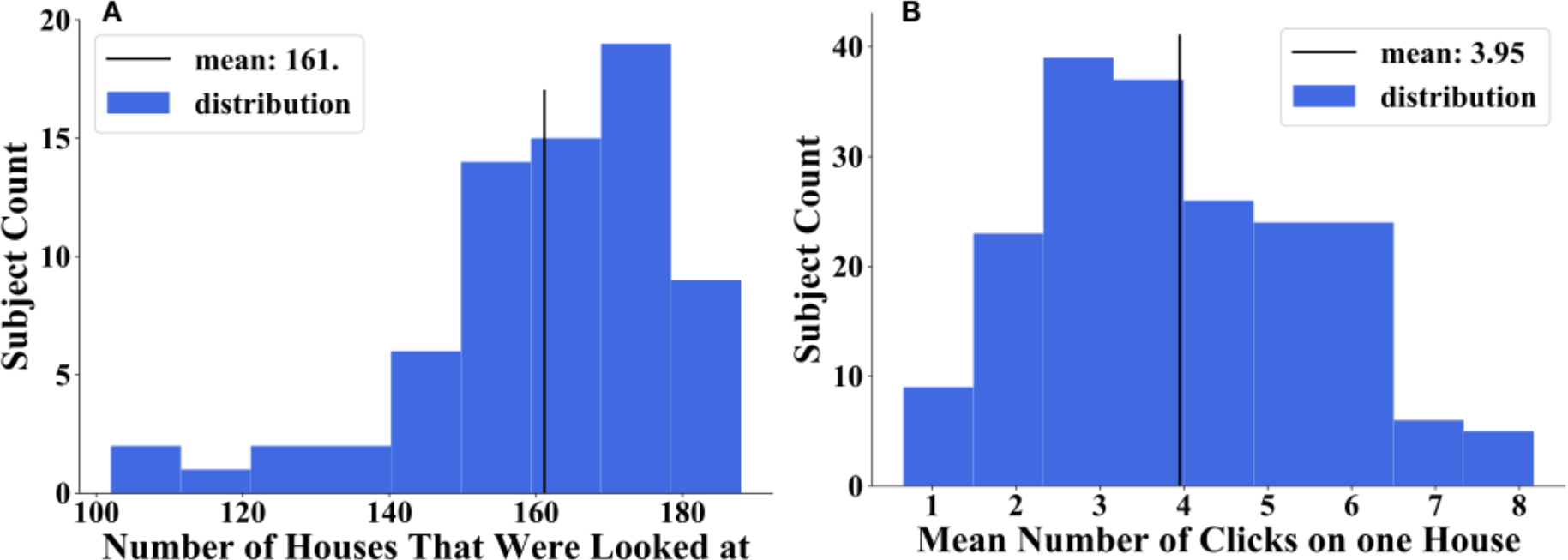
Distribution of number of houses that were looked at (A) and distribution of mean number of clicks on one house over all participants (B).

The visualization of the spatial distribution on the city map (Fig. 7) revealed a good coverage of clicked on and therefore viewed houses over the city map. Most houses were looked at by the majority of participants. The houses on the outer parts of the city were looked at somewhat more often than the more centrally located houses (Fig. 7A). Accordingly, houses on the outer parts of the city were more often clicked on, in contrast to centrally located houses (Fig. 7B). However, we did not find systematically neglected areas of the city. This shows that the spatial exploration with the interactive map covers the whole city with a bias to the outskirts of Seahaven.

**Fig. 7:**
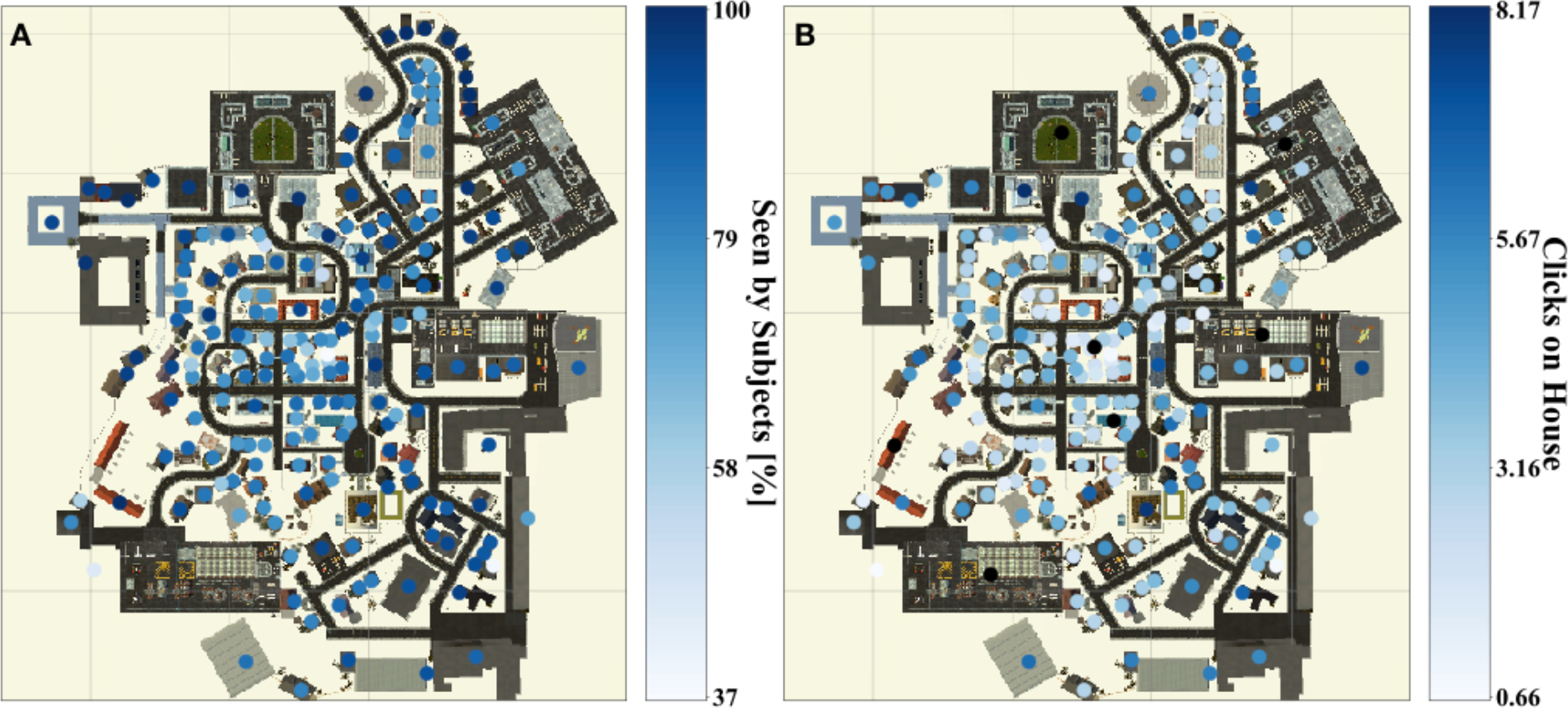
(A) Heat map depicting how many participants looked at a house and (B) how often participants clicked on houses on a gradient going from deep blue (most often clicked) to light blue (least often clicked). Black dots mark houses that could not be clicked on.

### Post-training Task Results

In this paper our main focus was to investigate the dependencies of performance in the post-training tests after one 30-minute training session. We first focused on a comparison of the different task and time conditions. Next, we considered the familiarity of houses, the distance between tested stimuli and the angular difference between tested stimuli as relevant factors. Additionally, we investigated the influence of subjectively rated abilities of spatial strategies using the FRS questionnaire (Münzer & Hölscher, 2011) on the task performance. As the last step of our analysis, we investigated the development of performance in the task and time conditions after repeated, i.e. up to three, training sessions.

#### Performance in different spatial task and decision time conditions after one session

We hypothesized that participants using an interactive city map for the exploration of the virtual city perform better with infinite time for a decision and thus for cognitive reasoning than in the spontaneous decision mode with 3 seconds response time. Furthermore, we assumed that participants training with a map would perform better in the absolute orientation task than in the relative orientation and pointing task.

Therefore, we calculated the fraction of correct answers per participant for each task and each time condition in a 3×2 design. This compared the spatial knowledge of absolute orientation, relative orientation and relative location of houses acquired during the exploration. We performed a repeated measure ANOVA with performance as the dependent variable, time (3sec, infinite) and task (absolute orientation, relative orientation and pointing) as repeated factor. We found a significant main effect for time (F(1,69) = 24,104, p < 0,001), but no significant main effect for task (F(2,138) = 1,693, p = 0,188) and no significant time*task interaction (F(2,138) = 1,393, p = 0,252). Our results revealed a performance in the 3 seconds condition slightly below 50% and significantly lower than in the infinite time condition. Effectively, this is a floor effect and a further investigation of the 3 seconds condition seemed to not be meaningful. Therefore, we calculated all following results taking into account only the infinite time condition. We observed mean performances in the absolute orientation task of 51,71%, in the relative orientation task of 54,72%, and in the pointing task of 53,41% (Fig. 8). However, we did not find a significant difference in task performances. In conclusion and in line with our hypothesis, we found that after 30 minutes of exploration of the virtual city with the interactive map, participants performed significantly better in the infinite time condition than under time constraints. This suggested that they needed time for cognitive reasoning to retrieve knowledge about the tested spatial properties and did not yet build up spontaneously retrievable knowledge.

**Fig. 8:**
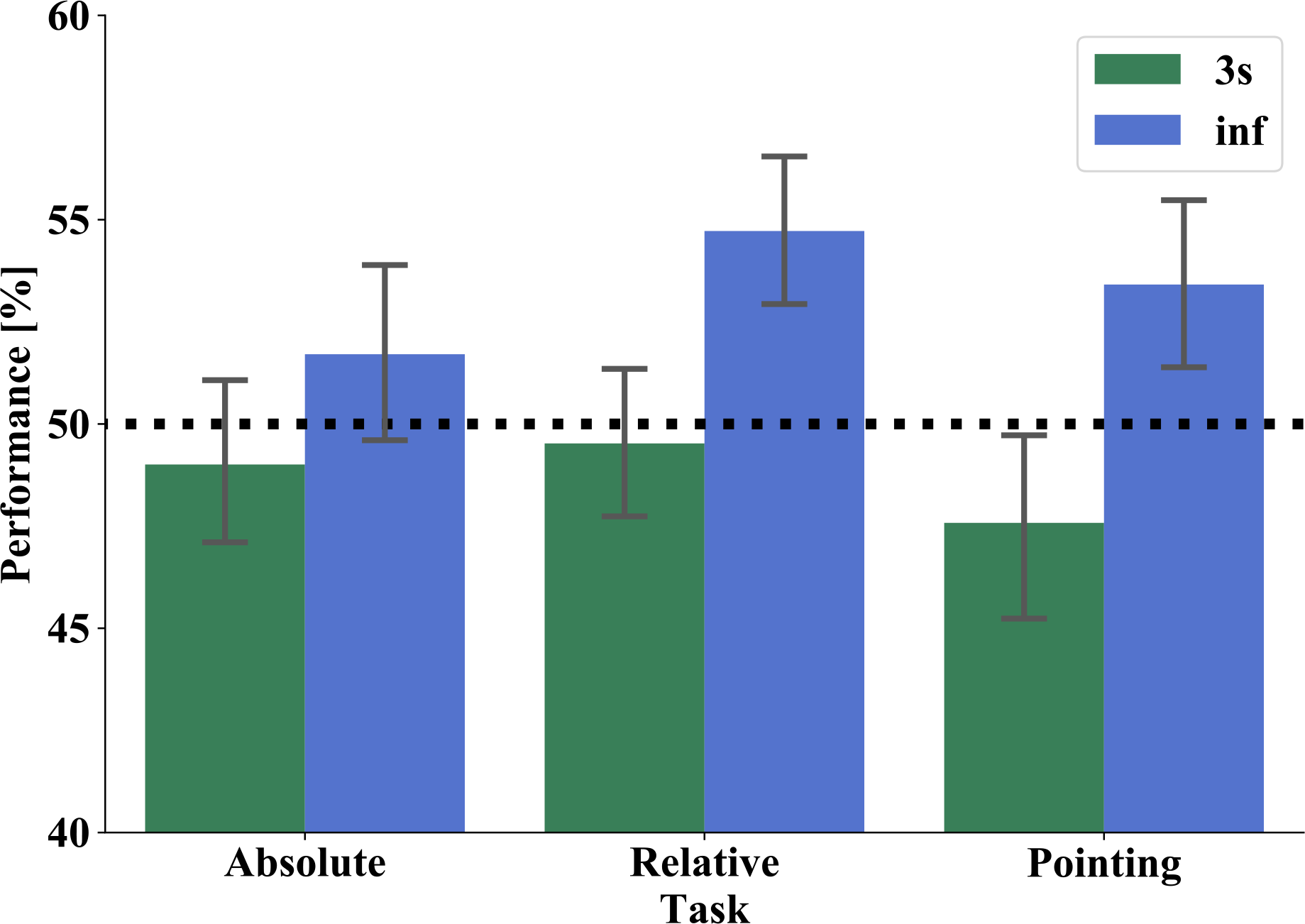
Performance of absolute orientation (left), relative orientation (middle) and pointing task (right) in 3 seconds (green) and infinite time (blue) condition. The bars show the mean performance of all subjects in both time response conditions. The black dashed line marks the chance level of 50%. The error bars indicate the standard deviation (SD).

#### Performance as a function of familiarity

We hypothesized that the better a house is known, the better the performance would be. Here, we used the number of clicks on a house as the familiarity measure. In other words: the more often a house was clicked on and viewed during the city exploration with the interactive map, the better we expected the performance to be. For the relative orientation task, we compared the clicks on the prime and the correct target house and for the pointing task the clicks on the prime and the target house in each trial. We then used the performance to the house with the lower click numbers for this trial. With this, we calculated for each participant the performance averaged over the trials containing houses with none, one, and two, and so on up to the maximum of clicks observed. The distribution of clicks by single participants ranged from 0 to 55 clicks on one house with a mean of 3.95 clicks. Because of the right skewed distribution we applied the natural logarithm to our data. With these we then performed a weighted linear regression (Fig. 9) separately for the three tasks. The weights are equal to the number of trials contributing to the mean performance for each number of clicks of each participant. For the absolute orientation task the results revealed a correlation with a slope of 0,0059 and an intercept of 0,51. Statistically this relation between logarithmic (ln) number of clicks and performance was not significant (F(1,603) = 0,2332, p = 0,629, R^2^ = 0,000). For the relative orientation tasks the results revealed a correlation with a slope of 0,0035 and an intercept of 0,54. This relation between logarithmic (ln) number of clicks and performance was not significant (F(1,469) = 0,074, p = 0,786, R^2^ = 0,000). For the pointing task the results revealed a correlation with a slope of 0,025 and an intercept of 0,51. This relation between logarithmic (ln) number of clicks and performance was not significant (F(1,481) = 3,617, p = 0,059, R^2^ = 0,007). In conclusion, we found no significant positive linear correlation of familiarity and performance in our tasks.

**Fig. 9:**
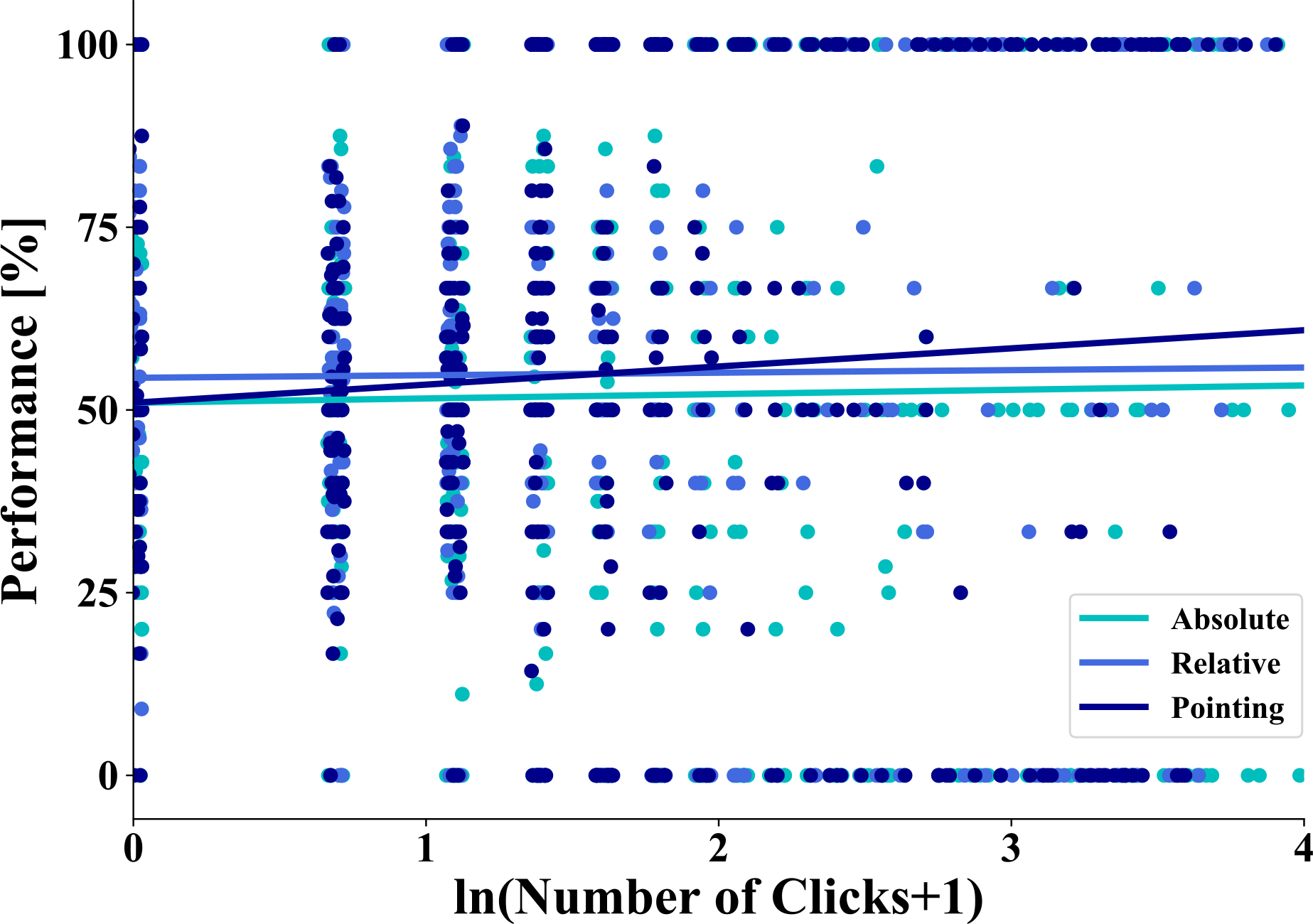
Linear regression between task performance separately for the three tasks and familiarity measured as the natural logarithm of the number of clicks on a house. One dot represents the average performance of all trials from one subject with a specific number of clicks on a house. The turquoise color depicts the data for the absolute task, the light blue for the relative task and the dark blue the pointing task.

**Fig. 10:**
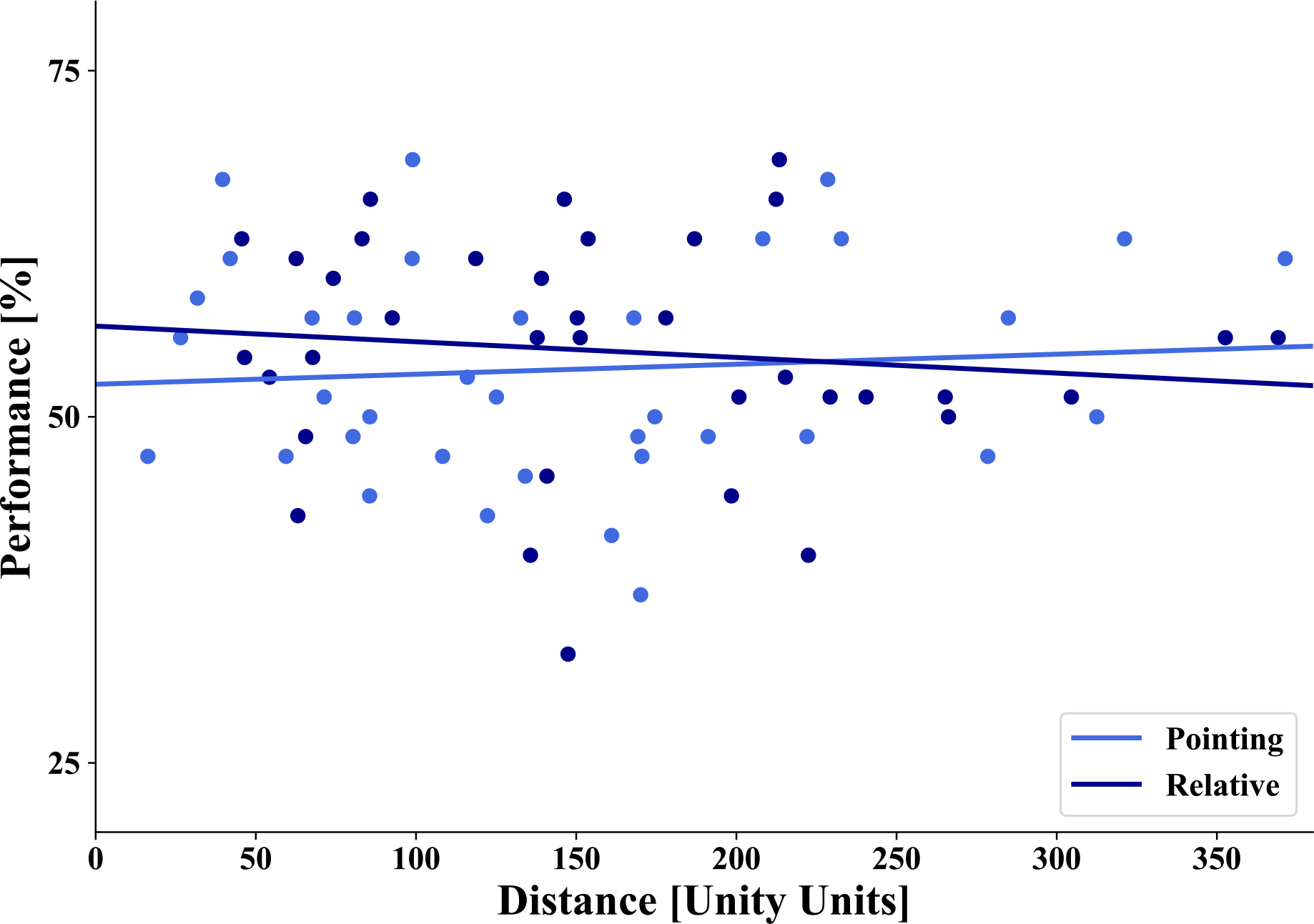
Linear regression of distance (abscissa) and performance (ordinate). The blue dots depict the combination of task and house averaged over subjects, dark blue for the relative orientation task and light blue for the pointing task. The straight lines depict the linear regression. We found no significant correlation between performance and distance.

### Performance as a function of distance

Participants learned the spatial dependencies of the virtual city with an interactive city map. Based on previous studies, we hypothesized that we would find no influence of distance between the houses on the performance (Loomis et al., 1993; Meilinger, Frankenstein, Watanabe, Bülthoff, & Hölscher, 2015). To estimate the influence of distance, we decided to only calculate the distance between two houses and not consider the complex triangular design in the relative task to perform comparable analysis to the pointing task. Therefore, we calculated the distance between prime and target house in the pointing task and between the prime and the correct target house in the relative orientation task. We investigated the distance up to 310 unity units in the relative task and up to 371 unity units in the pointing task representing the maximal distance of houses used in the respective task. We performed a linear regression analysis and found indeed no significant correlation between performance and distance for the relative orientation task (F(1,34) = 0,475, p = 0,495, R^2^ = 0,014) and the pointing task (F(1,34) = 0,236, p = 0,630, R^2^ = 0,007). Consistent with map learning, we found no distance effect when participants learned the city layout with the interactive map.

### Performance as a function of angular difference

Next we investigated whether after map learning the performance in the 2AFC task would depend on the size of the angular difference. In all tasks, participants had to compare two alternative choices that differed from each other in varying angular degrees in steps of 30° from 30° to 180°. We calculated the angular difference in the absolute orientation task between two arrows indicating cardinal directions on the stimuli and in the pointing task between two arrows pointing from a target house to a prime house. In the relative orientation task, we computed the angular difference between the orientations of two target houses. Due to the variations in orientation of the houses, the angular difference between these angles varied from a minimum of 30° in steps of 30° with a maximal deviation of +/− 5° in each step. We hypothesized that metric representations should make larger angular differences between the choices easier to discriminate and thus yielding a better performance. We observed the best performance at an angular difference between the options of 120° (Fig. 11). Furthermore, at 180° performance is close to chance level again. For statistical testing, we performed a one-way repeated measure ANOVA with performance as the dependent variable and angular difference categories (30°, 60°, 90°, 120°, 150°, 180°) as the repeated factor. However, in spite of the large number of subjects, we could not demonstrate a significant main effect for angular difference (F(5,345) = 1,5027, p = 0,1883). Thus, similarly to the lack of a distance effect between houses, we do not observe an effect of the angular difference on performance.

**Fig. 11:**
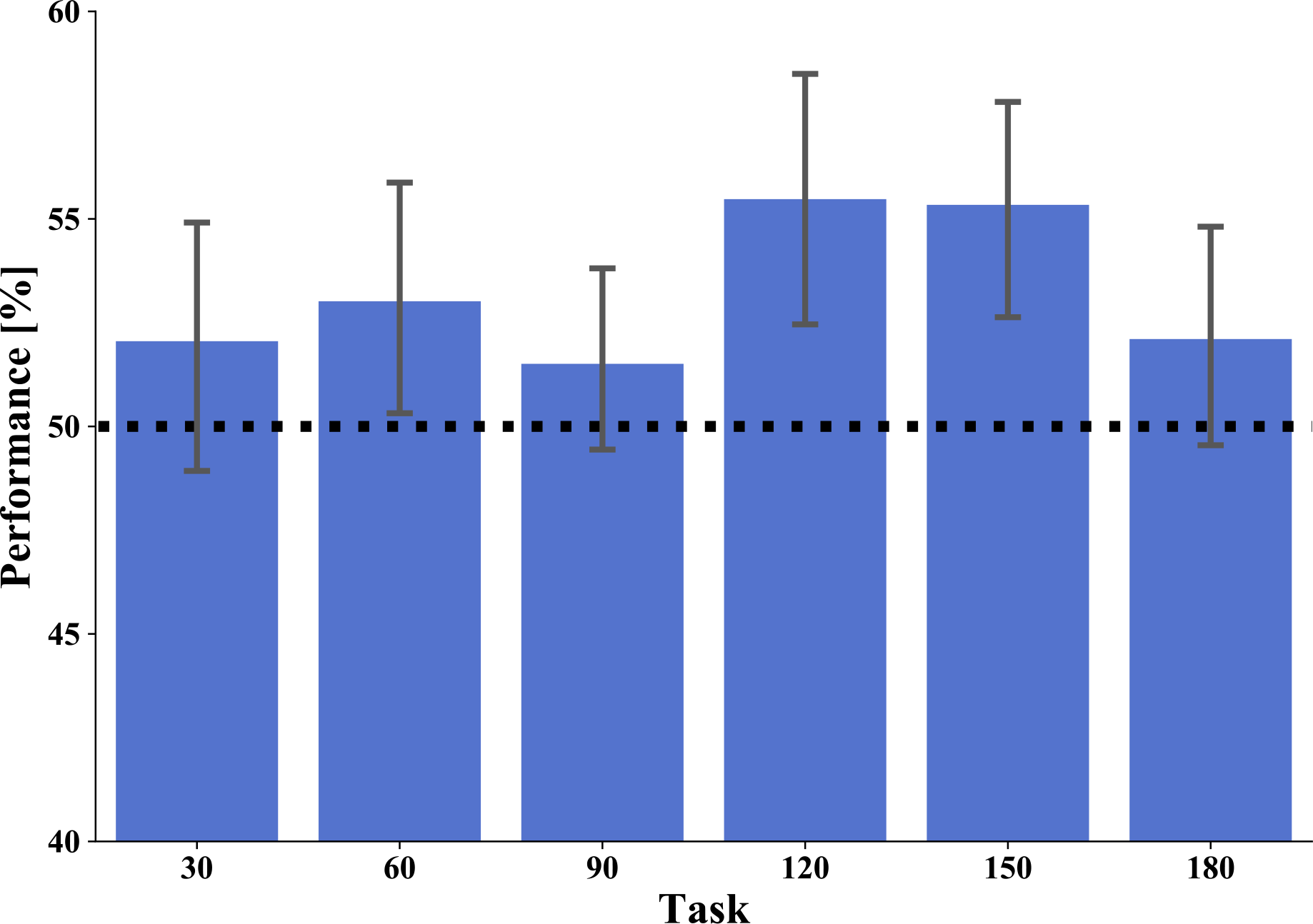
Performance in relation to angular difference between choices in the task stimuli over all tasks. Bars depict mean performance in 30°, 60°, 90°, 120°, 150° and 180° categories. Error bars depict standard deviation (SD). We see the best performance with an angular difference of 120° degrees but performance difference did not reach significance.

**Fig. 12:**
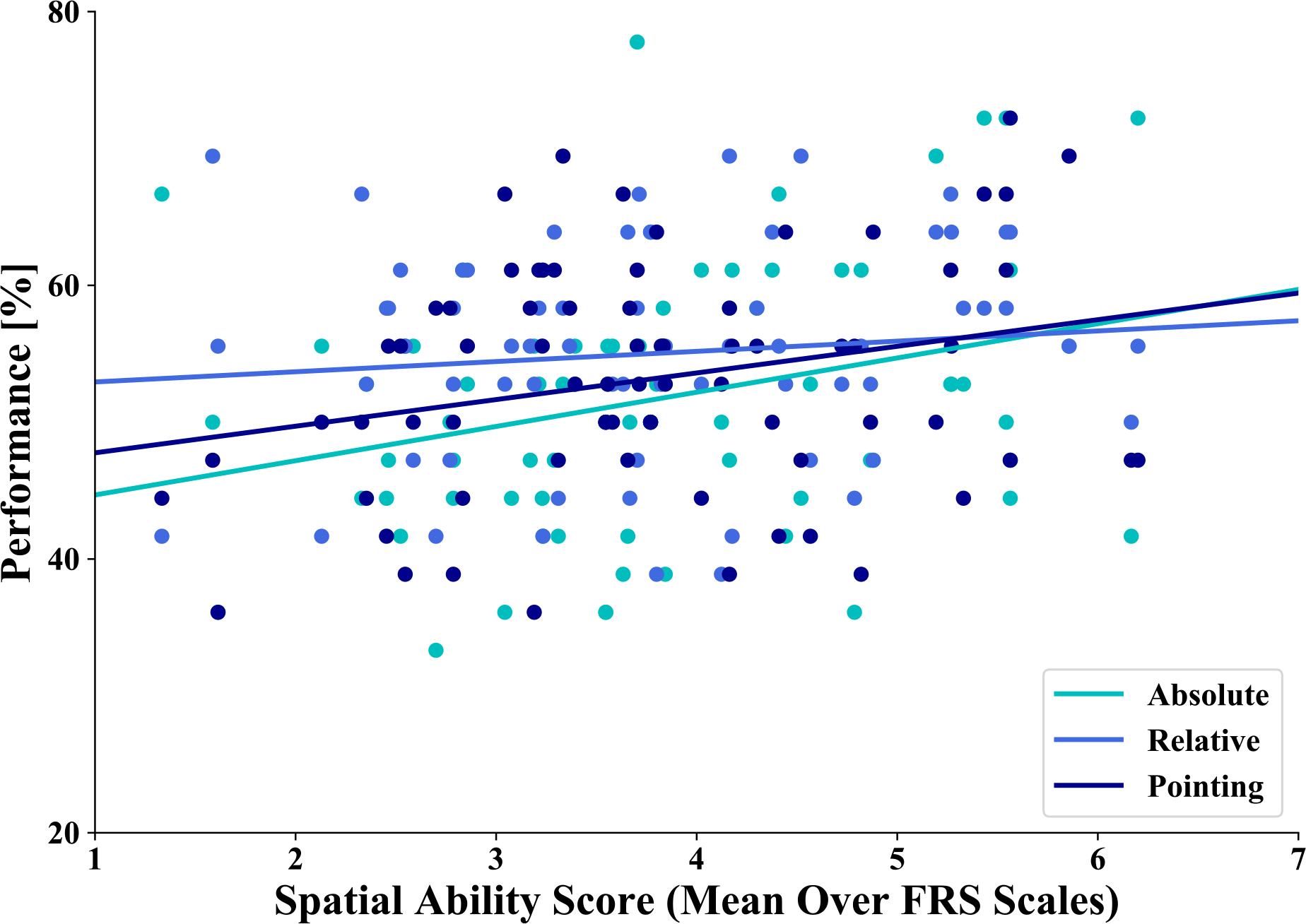
Linear regression between task performance (y-axis) and spatial ability score (x-axis). The straight lines depict the correlation: dark blue line correlation between performance in the absolute orientation task and the spatial ability score, light blue line correlation between the performance in the relative orientation task and the spatial ability score and the turquoise line correlation between the performance in the pointing task and the spatial ability score. The dots depict the single data points of the tasks in the respective colors.

**Fig. 13:**
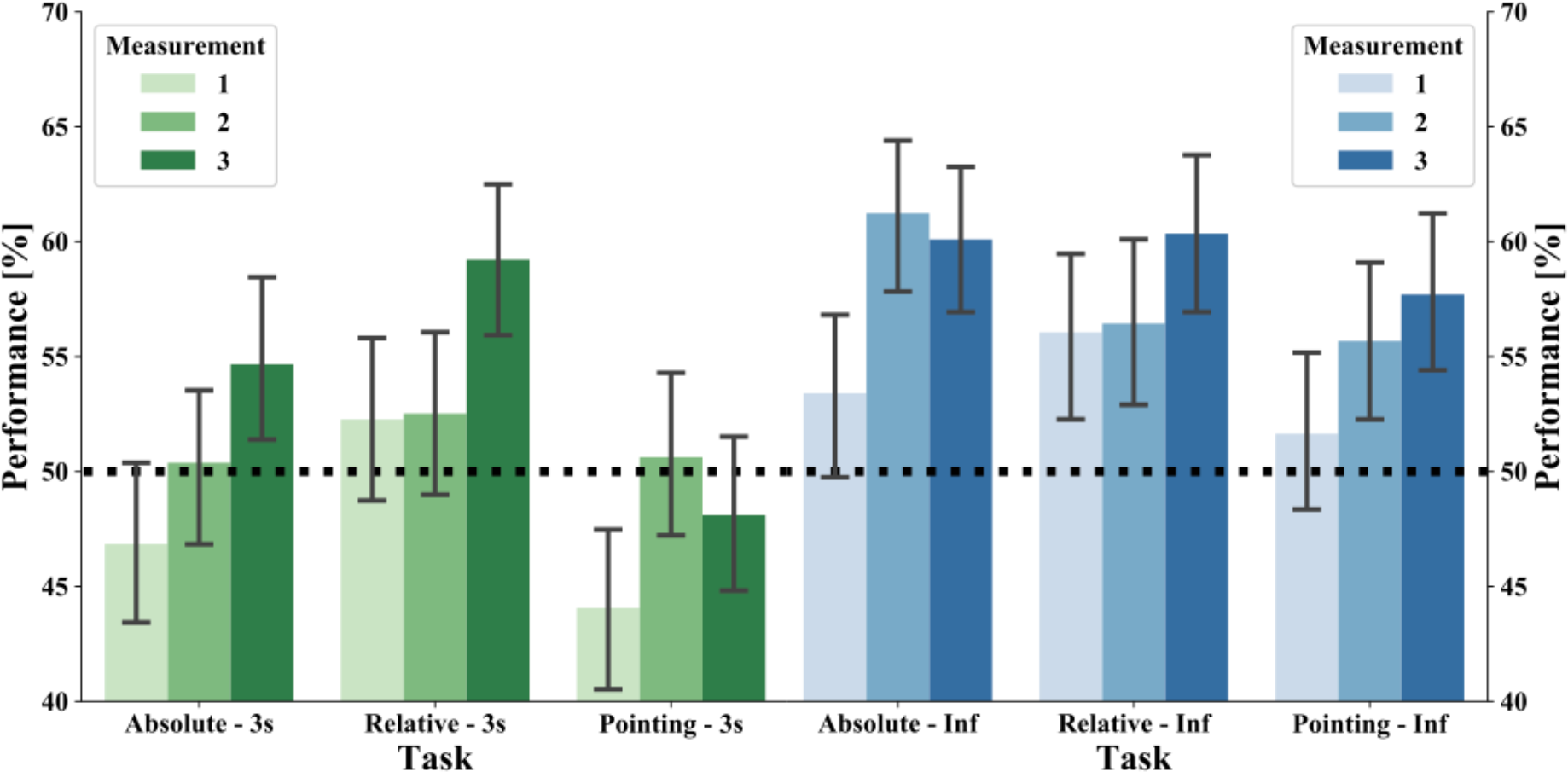
Performance of absolute orientation, relative orientation and pointing tasks in the infinite time condition with three repeated measurements. The green bars show the mean performance of all subjects in the 3 seconds response condition and the blue bars in the infinite time response condition. The light bars depict the mean performance on the first measurement day. The medium bars depict the mean performance on the second measurement day. The dark bars depict the mean performance on the third measurement day. The black dashed line marks the chance level of 50%. The error bars indicate the standard deviation (SD).

### Performance as a function of FRS scaling

To estimate subjectively rated abilities in spatial strategies and their influence on task performance, we used the FRS questionnaire. This questionnaire consists of three scales. Scale 1, the “global-egocentric orientation” scale captures general ability and egocentric strategies based on knowledge of routes and directions. Scale 2, the “survey” scale comprises of indicators of mental map formation. Scale 3, the “cardinal directions” scale measures the knowledge of cardinal directions. We calculated the Pearson correlation between the 3 FRS scales for our subjects (scale 1: M = 4,442, SD = 1,2; scale 2: M = 4,011, SD = 1,392; scale 3: M = 2,875, SD = 1,446). We found a strong positive significant pairwise correlation between all three scales of the FRS questionnaire (scale1/scale2: r = 0,577, p < 0,001; scale1/scale3: r = 0,418, p = 0,001; scale2/scale3: r = 0,528, p = 0,000). This indicates that participants who estimated themselves as good navigators did so in all spatial strategies and vice versa. As we were interested to investigate whether being a good or bad navigator would influence performance in the three tests, we decided to calculate a summary spatial ability score using the mean over all FRS scales. We then used this spatial ability score to calculate a correlation to the performance in the different tasks (absolute orientation, relative orientation and pointing task). We hypothesized that participants with a higher rated spatial ability would also perform better in our spatial tasks. We found a significant positive correlation between the spatial ability score and the performance in the absolute orientation task (R^2^ = 0,084, p = 0,013) and in the pointing task (R^2^ = 0,065, p = 0,03). Although average performance in the relative orientation task was the best of the three tasks, we found no correlation between the spatial ability score and this performance (R^2^ = 0,012, p = 0,364). Thus, subjects showed on average a high performance in the relative orientation task independent from their subjectively rated spatial ability. In contrast, the better a participant estimated his/her spatial ability, the better he/she performed in knowing the orientation towards cardinal north and house-to-house locations.

### Performance after repeated training sessions

To investigate, whether the task performance would increase with more time to explore the city, 22 (11 female) participants were trained and tested on three separate days within 2 weeks. Each time they trained for 30 minutes with the interactive map and afterwards performed the post-training tasks. As additional training could also improve the 3 seconds task conditions, we calculated a repeated measure ANOVA with performance as the dependent variable and task (absolute orientation, relative orientation, pointing task), time (3 seconds, infinite) and measurement day (1, 2, 3) as repeated factors in a 3×2×3 design. We found a significant main effect for task (F(2,42) = 5,630, p = 0,007). The grand average performance in the tasks was 54,3%, 55,9%, and 51,6% for absolute, relative and pointing, respectively. Furthermore, we observed a significant main effect for time (F(1,21 = 26,853, p < 0,001). The grand average performance in the 3 seconds and infinite time conditions was 51,2%, and 56,6%, respectively. Finally, we observed a main effect for measurement day (F(2,42) = 9,022, p < 0,001). The grand average performance was 50,7%, 54,4%, and 56,7% for first, second and third measurement day, respectively. We found no significant two way interactions (task*time: F(2,42) = 2,430, p = 0,1; tasks*measurement: F(4,84) = 2,108, p = 0,087; time*measurement day: F(2,42) = 0,176, p = 0,839) and no significant three way interaction (F(4,84) = 1,305, p=0,275). In summary, our data suggest that with more time to explore the city with the interactive map, participants improve their performance in the spatial tasks, revealing a better performance with time for cognitive reasoning and a better performance in the absolute and relative orientation tasks than in the pointing task.

## Discussion

In the present study, we investigated learning of spatial properties after exploration of a large-scale virtual city with an interactive city map after single and repeated 30-minute training sessions. After each session, we performed three tasks adapted from a previous study (König et al., 2017) to measure spatial knowledge of houses’ orientation towards cardinal north (absolute orientation), orientation of two houses to each other (relative orientation) and the location of two houses to each other (pointing task). Our results revealed that one-time training with the interactive city map was not sufficient enough to acquire spontaneous spatial knowledge, but time for cognitive reasoning improved the performance significantly. With repeated training sessions, performance increased significantly and, additionally, significant task differences were revealed. Specifically, performance in the pointing task was lower than performance in the other two tasks. We found a positive correlation of the subjective self-assessment of spatial strategies to the performance in the absolute orientation and pointing task but no correlation to the relative orientation task. Our results revealed no correlation of familiarity measured in how often houses were looked at onto the performance level. Training with our map did not result in a dependence of performance on the distance between houses (absence of distance effect). Further, we did not observe a dependence of angular difference between stimuli on performance (absence of angular difference effect). We argue that this pattern of results is compatible with an interpretation that after interactive map training information is memorized on an abstract level and not truly tapping into resources of spatial cognition from an embodied perspective.

In the present study, we observed a performance level only slightly above chance after 30 minutes exploration with the interactive city map and time for cognitive reasoning in the retrieval tasks. Our task design is in line with previous studies in which participants also performed 20-30 minutes training with the environment, which was followed by a test phase shortly afterwards (Nathan Greenauer & Waller, 2008; Marchette et al., 2011; McNamara et al., 2003; Mou & McNamara, 2002; Mou, Zhang, & McNamara, 2009; Shelton & McNamara, 2001). However, it has to be considered that in relation to previous studies the extent and complexity of our virtual city was large and the training time of half an hour was relatively short for a city map of that spatial extent with a lot of material to be absorbed and memorized. This time limitation was motivated by the training, which was performed in actually navigating through the virtual city. Here, extended training beyond 30 minutes would lead to more frequent and more severe motion sickness. We therefore decided on a training time of 30 minutes for one session equally for all three experiments using the virtual city including the exploration with the interactive map. To counterbalance possible effects of training time limitation, we are currently performing repeated training sessions, reporting here first results on these sessions. Even though, the one-time training was sufficient so that the vast majority of houses were looked at and most of the houses could be viewed several times. In contrast, getting to know a real world environment takes place over much longer time scales, thus allowing spatial knowledge to evolve over a much longer time span. Increased familiarity with an environment improves spatial knowledge (Burte & Montello, 2017; Ishikawa & Montello, 2006; Lindberg & Gärling, 1983; Montello, 1998). In line with these findings, the performance in the present study improved significantly. After three training sessions the average performance level of 56,7% is sizable in comparison to the previous study by König et al. (2017). There, subjects had experience with a real world city for at least a year and performed on average at 59,1%. Concluding, our training and task design is demanding, but given the restricted time to explore our virtual city the performance level is reasonable and increased time to explore the virtual city improves knowledge of spatial properties.

One objective of the current study was to test previous reports (König et al., 2017) in a highly controlled laboratory setup. That paper investigated spatial knowledge after at least one-year experience in the hometown. They reported a better knowledge of relative house-to-house orientation and location than of absolute orientation towards north with restricted response time, and a reversal of that relation with infinite time available. In that paper, the best performance was found in pointing from one house to another house. Those results were argued to be in line with an action oriented embodied approach to spatial cognition. In the present study, analyzing the training with the interactive map of our virtual city revealed that participants looked at the majority of houses and viewed a house several times. The performance in the spatial tasks in the 3 seconds response condition, however, was at chance level. Time for cognitive reasoning was needed to improve the task performance. It appears that the available training time was not yet sufficient to build intuitively available knowledge or reach discernable performance differences between the tasks. Indeed, task performance improved significantly over repeated training sessions revealing differences between task and time conditions, indicating that with enough training time, differences in spatial learning become noticeable. In contrast to the previous study in which performance in the pointing task was best, in the present study the subjects training repeated sessions revealed the worst performance in the pointing task. This indicates that the acquisition of spatial knowledge by an interactive 2D map taps on different mechanisms than spatial learning in a real world city. Taking the number of clicks on a house as an objective familiarity measure, we found no significant correlation between performance and familiarity after a single training session. In the interpretation of this surprising result we have to consider, that the number of clicks on a house is not an independent, but a dependent variable. That is, subjects might find specific houses more difficult to memorize and, therefore, click more often onto these houses. Thus, the subjectively perceived familiarity after training might diverge from an objectively defined familiarity based on behavior during training. König et al. (2017) additionally investigated spatial knowledge of streets’ orientation and found that streets were coded with spontaneous retrieval in relation to cardinal north. As the layout of our virtual city did not allow for adequate numbers of streets, we did not test this aspect in the current paper. Investigating the effect of angular differences between tested stimuli choices, the previous study found better performance with larger angular differences, thus a positive angular difference effect. In contrast, after a single session of spatial learning with our interactive city map, we found no significant angular difference. This lack of angular difference effect would be explainable if angular differences could be coded in a categorical form after map learning, allowing a same/different judgment, but not a metric comparison. Again, it is an indication that the acquisition of spatial knowledge by an interactive 2D map is qualitatively different to spatial learning in the real world. Taken together, the differences of our current results to the previous study suggest that learning with an interactive map supports learning of spatial properties differently than getting to know an environment by the embodied perspective of everyday navigation.

Though, previous research found large individual differences in spatial knowledge acquisition (Burte & Montello, 2017; Hegarty, Richardson, Montello, Lovelace, & Subbiah, 2002; Hegarty et al., 2018; Ishikawa & Montello, 2006; Montello, 1998; Wolbers & Hegarty, 2010). Ishikawa and Montello (Ishikawa & Montello, 2006) found especially large differences in learning survey knowledge, which was positively correlated to a subjective report of the Santa Barbara Sense-of-Direction (Burte & Montello, 2017; Hegarty et al., 2002). The Santa Barbara Sense-of-Direction was especially correlated with pointing to locations in a known environment (Hegarty et al., 2002; Sholl, 1988), spatial knowledge in a newly learned environment (Hegarty, Montello, Richardson, Ishikawa, & Lovelace, 2006), the allocentric heading-recall task (Burte & Hegarty, 2012; Sholl, Kenny, & DellaPorta, 2006) and recently also to the ability to point to north (Brunyé et al., 2015; Burte & Montello, 2017). As indicated by Wolbers and Hegarty (Wolbers & Hegarty, 2010) being a successful navigator might be especially determined by choosing the right spatial strategy for a spatial task. In our study, we investigated spatial abilities using the “Fragebogen Räumliche Strategien” FRS (translated: Questionnaire of Spatial Strategies). This questionnaire (Münzer & Hölscher, 2011) measures spatial orientation strategies and was validated to predict spatial learning in real environments. In our study, we wanted to evaluate whether subjective reports of spatial learning in real environments would also predict learning of spatial properties with a map of a virtual city. Investigating the three FRS scales revealed a strong correlation between these, indicating that a participant who judged his spatial abilities as being good did so in all three scales and vise versa. As we were interested in the influence of good or bad navigators on the performance in our tasks, we calculated a spatial ability score comprised of the three FRS scales combined. In line with previous studies, we found a positive correlation between this spatial ability score and the performance in our pointing task testing the knowledge of spatial relations between house locations and a positive correlation to the performance in our absolute orientation task testing knowledge of the orientation of houses towards cardinal north. We did not find a correlation between the combined FRS scales and the performance in the relative orientation task testing knowledge of relative orientation between houses, although the performance in that task was higher than performance in the pointing task. This, however, violates expectations of a positive correlation i.e. that good navigators should learn spatial tasks well and reach a higher performance than bad navigators. A possible explanation is that acquisition of information for the relative orientation task by exploration of a 2D interactive map does not tap on resources of every-day spatial cognition, but is acquired differently. In conclusion, real world differences of spatial abilities measured in a subjective self-report might also predict the learning of spatial properties with an interactive city map of a virtual city.

Another factor that might influence spatial knowledge acquisition is whether it is achieved incidentally or only with intention. Burte & Montello (2017) investigated how the sense of direction and learning intentionality influences spatial knowledge acquisition. They found that participants with a good self-reported sense of direction were more accurate in acquiring spatial knowledge by direct experience while intentionality had no learning effect. Regarding intentional learning, we decided to instruct and familiarize all participants in a pre training task before the exploration with the map about the post-training task design. A pilot study had revealed that some participants paid attention to not spatially relevant features, e.g. details of house design during the training phase. Thus, we made sure that all participants were able to intentionally pay attention to relevant spatial features. Furthermore, when performing several sessions, participants already know what is to come after the first session. Thus, the pre-task helps to make the comparison of first and later sessions valid, even though we thus cannot investigate possible effects of intentional learning.

Getting to know a new environment with the aid of a map is supposed to support the acquisition of survey knowledge especially (Frankenstein et al., 2012; Meilinger et al., 2013; Meilinger et al., 2015; Taylor et al., 1999; Thorndyke & Hayes-Roth, 1982) and the acquisition of a global reference frame (Frankenstein et al., 2012; Meilinger et al., 2015; Richardson, Montello, & Hegarty, 1999). Survey tasks can also be solved using global reference frames that are learned from experience leading to a decrease in performance with greater distance (distance effect) (Loomis et al., 1993; Meilinger et al., 2015). Using a global reference derived from a map is not supposed to yield a distance effect (Frankenstein et al., 2012) because a map directly provides an overview giving distances and locations as well as spatial orientation in proportion to the depicted environment. In line with previous research in the presented study, we did not find a distance effect on performance in our tasks after learning with our interactive map. In close analogy to the lack of distance effect, we did not find an effect of angular differences after learning with the map. These results suggest that participants acquired the knowledge of a global reference frame not only including the information of distance, but also of angular information while training with the city map.

Reference frames are important to organize learning and memory of spatial properties in small and large environments (Greenauer & Waller, 2010). For actual navigation in the environment, allocentric spatial relations are recalled and are combined with present egocentric information (Kelly & McNamara, 2008; Mou et al., 2004; Riecke et al., 2007). Beside egocentric and allocentric reference frames intrinsic object-to-object relation is an important reference to learn and memorize spatial relations (Mou & McNamara, 2002). Egocentric spatial cues, which are combined with allocentric environmental cues, are supposed to be the main cues for building an intrinsic reference frame (McNamara, 2002). However, Mou and McNamara (Mou & McNamara, 2002) found that objects’ locations were remembered in relation to a salient intrinsic reference frame defined by the layout of the environment even when this was not aligned to the egocentric experience. In our study, the setup of the map training makes use of an allocentric reference frame related to absolute north (up on the map) intuitive as our interactive map depicts a conventional two-dimensional city map of our virtual city.

The task design in the current study was based on front on screenshots of houses taken in our virtual city from a pedestrian perspective to replicate the study of König et al. (2017) in a virtual city. Therefore, learning the city layout with a city map required an interactive feature that combined the map with front on views of the houses. Here, we decided on adding a feature to the city map that enabled to click on single houses from a top view. By clicking on a house, the respective house front taken as a screenshot was displayed on the two screens beside the map. This lead to a necessary change of perspective from viewing the map from a bird’s eye view to watching the house fronts from a horizontal perspective. Previous research found that recalling spatial information in a different orientation or perspective than it was learned and stored yields, costs in terms of increased errors or delays (McNamara, Sluzenski, & Rump, 2008; Meilinger et al., 2013; Shelton & McNamara, 2004).

In contrast to a perspective switch between acquiring and retrieving spatial properties, here participants had to switch the perspective during the training with the interactive map. However, this does not allow for a three-dimensional exploration of the city like in VR and therefore we are not providing participants with an embodied experience of a three-dimensional map. Comparing learning with a 2D and a 3D city map, researchers found a better performance in spatial tasks after learning with a 2D map, even with participants who were experienced in virtual reality games (Oulasvirta et al., 2009). Our results did not show a better performance in knowing the orientation of a single house towards cardinal north than in the other tasks after one training session. After repeated training sessions, we found a better performance in the absolute, as well as the relative orientation task compared to the pointing task. This indicates that with more training time metric spatial information, which is directly available from a map resulted in better performance. Taken together, our results suggest that after training with an interactive city map, spatial information is memorized on an abstract level, mirroring the metric information given by the map and not truly tapping into resources of spatial cognition from an embodied perspective.

## Conclusion

In the present study, we investigated the learning of spatial properties in the exploration of a large-scale virtual city with an interactive city map in a controlled set up. Our results revealed that our design was valid to explore most of the houses a few times within a one-time training session. They suggest that repeated trainings are necessary to improve performance and reveal task differences with a better performance in the absolute and relative orientation than in the pointing task. They further suggest that participants acquired a global reference frame when learning with the map indicated by a lack of distance and a lack of angular difference effects. These results seem to indicate that solely learning with a map, spatial information is acquired and memorized on an abstract level thus in contrast with an action-oriented approach. The next step in future research is to investigate learning of spatial properties while actually exploring the virtual city in VR. This offers the possibility of comparing acquired spatial knowledge with controlled learning either with the interactive city map or directly navigating in VR, giving insight into differently learned spatial properties.

## Acknowledgements

We gratefully acknowledge financial support by the European Commission (H2020 FETPROACT-2014, SEP-21014273, socSMCs, ID: 641321, PK), by Deutsche Forschungsgemeinschaft (DFG) (Research Training Group on Situated Cognition (RTG 2185), NK) and the Open Access Publishing Fund of Osnabrück University.

## Author Contribution Statement

SK designed the study, supervised and managed the study, analyzed the data, wrote and revised the paper.

VK built the virtual city, wrote scripts, analyzed the data and revised the paper.

DN built the map, wrote the code for the map and revised the paper.

LD measured participants, wrote scripts and analyzed data.

NK measured participants and revised the paper.

PK designed the study, supervised the study, and revised the paper.

